# Sleep Dependent Changes of Lactate Concentration in Human Brain

**DOI:** 10.1101/2021.12.05.471196

**Authors:** Selda Yildiz, Miranda M. Lim, Manoj K. Sammi, Katherine Powers, Charles F. Murchison, Jeffrey J. Iliff, William D. Rooney

## Abstract

Lactate is an important cellular metabolite that is present at high concentrations in the brain, both within cells and in the extracellular space between cells. Small animal studies demonstrated high extracellular concentrations of lactate during wakefulness with reductions during sleep and/or anesthesia with a recent study suggesting the glymphatic activity as the mechanism for the reduction of lactate concentrations. We have recently developed a rigorous non-invasive imaging approach combining simultaneous magnetic resonance spectroscopy (MRS) and polysomnography (PSG) measurements, and here, we present the first *in-vivo* evaluation of brain lactate levels during sleep-wake cycles in young healthy humans. First, we collected single voxel proton MRS (1H-MRS) data at the posterior cingulate with high temporal resolution (every 7.5 sec), and simultaneously recorded PSG data while temporally registering with ^1^H-MRS time-series. Second, we evaluated PSG data in 30 s epochs, and classified into four stages Wake (W), Non-REM sleep stage 1 (N1), Non-REM sleep stage 2 (N2), and Non-REM sleep stage 3 (N3). Third, we determined lactate signal intensity from each 7.5-s spectrum, normalized to corresponding water signal, and averaged over 30-s for each PSG epoch. In examinations of nine healthy participants (four females, five males; mean age 24.2 (±2; SD) years; age range: 21-27 years) undergoing up to 3-hour simultaneous MRS/PSG recordings, we observed a group mean reduction of [4.9 ± 4.9] % in N1, [10.4 ± 5.2] % in N2, and [24.0 ± 5.8] % in N3 when compared to W. Our finding is consistent with more than 70 years of invasive lactate measurements from small animal studies. In addition, reduced brain lactate was accompanied by a significant reduction the apparent diffusion coefficient of brain lactate. Taken together, these findings are consistent with the loss of lactate from the extracellular space during sleep while suggesting lactate metabolism is altered and/or lactate clearance via glymphatic exchange is increased during sleep.

**Significance Statement:** This study describes a non-invasive magnetic resonance spectroscopy/polysomnography technique that allows rigorous measurement of brain metabolite levels together with simultaneous characterization of brain arousal state as either wakeful or one of the several sleep states. The results provide the first *in-vivo* demonstration of reductions in brain lactate concentration and diffusivity during sleep versus wakefulness in young healthy human brain. These findings are consistent with invasive small-animal studies showing the loss of extracellular lactate during sleep, and support the notion of altered lactate metabolism and/or increased glymphatic activity in sleeping human brain.

## Introduction

While fundamental aspects of sleep physiology remain incompletely understood, it is clear that sleep is crucial to sustain brain health across the lifespan. Acute sleep deprivation results in impaired performance on attention and working memory tasks while chronic sleep disruption has profound impacts on overall daytime mood, motor, creative thinking, and cognitive performance. ^1–5^ Emerging epidemiological evidence further demonstrates the significance of sleep as sleep disruption is shown to be associated both with dementia and Alzheimer’s diagnosis^6,7^ as well as the development of amyloid plaques prior to the onset of clinical symptoms. ^8,9^ Despite the clear importance of sleep for acute and long-term brain health, underlying mechanisms associated with this increased risk remain poorly understood.

The potential metabolic benefits and restorative aspects of sleep have long been discussed, including reduced catabolic activity and increased anabolic activity.^10,11^ Lactate is an important cerebral metabolite that is present at high concentrations both within cells and also in the extracellular space between cells, the interstitial space. Cellular mechanisms underlying changes in brain lactate concentrations have been investigated in animal studies for more than 70 years,^12^ demonstrating sharp reductions in lactate concentration (12-35%) during sleep or anesthesia relative to wakefulness.^12–17^ In human brain, metabolic activity associated with cellular energy production is reduced during sleep with individual sleep stages associated with differing regional effects on brain metabolism,^18^ although gene expression indicating anabolic processes such as macromolecular synthesis is strongly increased.^19^

Along with its known dynamic profile in cerebral metabolism, lactate levels have also been used as a surrogate marker for clearance of brain interstitial fluid (ISF). ^13^ Such markers are relevant for the assessment of glymphatic activity, a physiology which supports the interchange of extracellular space fluids; cerebrospinal fluid (CSF) and ISF along perivascular pathways and facilitates the clearance of solutes and metabolic wastes from the brain interstitium.^20,21^ In rodent models, sleep and arousal state regulate glymphatic activity, as CSF tracer influx and interstitial solute clearance are increased during both sleep and some anesthesia conditions compared to the waking states.^22^

To our knowledge, alterations in human cerebral lactate concentrations across sleep-wake cycles, including different sleep stages, have not been reported. Given the important role of lactate in cerebral metabolism and its potential as a surrogate of glymphatic physiology, improved understanding of brain lactate dynamics across sleep-wake cycles is important for assessment of fundamental sleep-associated physiology and metabolism. Our primary goals in this study were to investigate cerebral lactate concentration across natural sleep-wake cycles and evaluate the potential of lactate as a sleep biomarker in human brain. To realize these goals, we successfully developed and applied a novel non-invasive experimental methodology combining rigorous and simultaneous magnetic resonance spectroscopy (MRS) and polysomnography (PSG) measurements in young healthy humans.

## Materials and Methods

The study was performed at a single site with all procedures approved by the Institutional Review Board of the Oregon Health & Science University (OHSU). All subjects provided verbal and written informed consent prior to study procedures.

### Study Subjects

To examine human brain lactate levels across sleep-wake cycles, we recruited young healthy subjects for the sleep-wake study experiments in a 3T MRI instrument. We recruited healthy subjects from the Portland metropolitan area using OHSU’s study participation opportunities website, Oregon Center for Clinical and Translational Research Institute (OCTRI) research match for recruitment, and flyers throughout the OHSU campus and communities in Portland. **Fig. 1** shows the study flow chart with the details of subject recruitment and enrollment procedures including a comprehensive screening with a list of inclusion and exclusion criteria. Of the 55 participants contacted for the study, 36 were phone screened, 19 were eligible for an initial visit for detailed screening, 14 were enrolled for sleep-wake experiments, and 9 subjects (four females, five males; mean age 24.2 ± 2 (SD) years; age range: 21-27 years) successfully completed the study and were included in the final data analysis.

**Figure 1.**
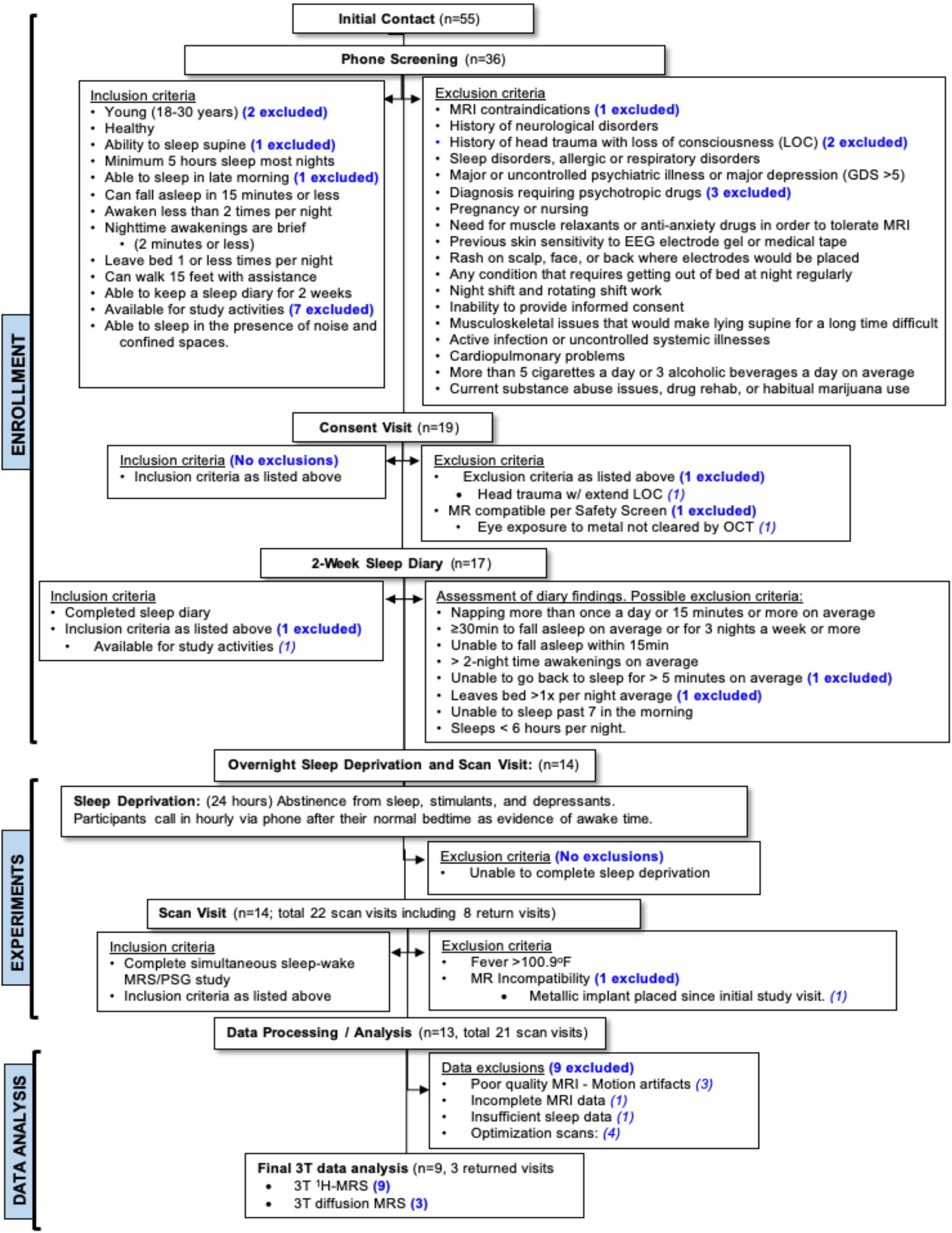
Study flow chart

The MRI environment is not conducive of restful sleep, as it is a confined space which can be acoustically noisy, with subjects restricted to the supine position, and passive restraints to reduce motion can increase discomfort. To increase the likelihood of sleep within the MRI environment, i) a 2-week sleep diary screening was utilized to enroll participants that are likely to sleep in the MRI environment, and ii) all enrolled subjects were sleep-deprived overnight, and were instructed to refrain from sleep, daytime napping, and caffeine and alcohol consumption for 24-hours prior to the study. The MRS study visits took place between 5:45 am-11:00 am after an overnight sleep deprivation, and included 1.5-hour subject PSG head cap preparation and up to 3-hour simultaneous MRI/MRS and PSG sessions. To ensure overnight sleep-deprivation, subjects were instructed to call-in every ~60 min and record a message throughout the night until their scheduled transportation arrived.

All subjects underwent sleep-wake experiments consisting of simultaneous recordings of MRS and PSG measurements in a 3T Siemens Prisma MRI instrument (*Siemens Healthineers, Erlangen, Germany*) using a 64-channel head/neck coil. PSG data were recorded using an MR compatible system (*Brain Products Inc, GmbH, Munich, Germany*) which involved (1) a customized 15-channel head-cap with 12 electroencephalogram (EEG) electrodes, two electrooculography (EOG) electrodes, and one drop-down electrocardiogram (ECG) electrode positioned on the back of the subject, (2) a respiration belt (*Respiration Belt MR*) placed on the chest to measure breathing and (3) a 3D acceleration sensor (*3D Acceleration Sensor MR*) positioned on subject’s left calf muscle to measure limb movements. Note that per vendor compatibility with MRI, PSG measurements did not include electromyogram (EMG) data collection.

### Subject PSG preparation

Prior to positioning in the MR instrument, each subject was prepared for PSG recording by study team members. Two different sized head-caps (*Brain Products Inc, GmbH, Munich, Germany*), with circumference of 56 cm or 58 cm, were used with the cap selected to best fit the subject’s head size. After placement of the head-cap, all electrodes were filled with an electrolyte gel (*Abralyt HiCI, Easycap GmbH*) to ensure consistent low electrode impedance throughout the experiment. Electrode impedances were displayed on BrainVision Recorder software: EEG and EOG: < 3kΩ and ECG: < 20 kΩ. A physiologic calibration test was performed to verify that all data channels delivered a physiologic signal before and after positioning the subject in the MR unit. Electrode impedance checks were performed every 30-60 min throughout the study and adjusted as needed to ensure subject safety.

### Subject positioning in the MR instrument

Prior to data acquisition, subjects were instructed to lie still in the supine position during the entire MR-PSG data acquisition. Subjects were provided with earplugs and headphones to attenuate acoustic noise and ease communication with the study team. Sufficient padding was provided to secure the head to reduce motion artifacts and prevent discomfort from occipitally-placed electrodes. Subjects were also provided with a bolster placed under the knees, foam pads placed under the elbows, and blankets for warmth during the MR-PSG session. The lights in the magnet bore and magnet room were off for the duration of the MRS study.

### Data Acquisition and Processing

#### Polysomnography

PSG measurements were continuously recorded with a sampling frequency of 5 kHz using *BrainVision Recorder* software, and temporally registered with ^1^H-MRS time-series using time-stamps - generated by the MR instrument and recorded with PSG. Due to coupling between the PSG system and the MR scanner during its normal operation, the raw sleep EEG, EOG, and ECG signals ranged from slightly to markedly obscured. In order to visualize real-time polysomnograms, *BrainVision RecView* software was used during the study, and *BrainVision Analyzer2* software for the post data-processing. In both cases, data processing included: 1) removing MRI coupled signals and cardioballistic artifacts, 2) down-sampling the data by a factor of 5 with a low-pass frequency of 100 Hz, 3) further band-passing the filtered signals as follows: EEG and EOG: [0.3 −20 Hz], EEG: [0.3-10Hz], respiration: [0.05-10 Hz], acceleration: [10-70 Hz] for visual inspection of each physiologic component.

Post-processed PSG data were evaluated in 30 s epochs. Sleep stages were visually scored by three experts based on modified criteria from the American Academy of Sleep Medicine (AASM), substituting EMG with a 3D accelerometer^23,24^ and classified into one of four stages by at least two independent experimenters (**Fig. 2B**): i. Wake (W), ii. Non-REM sleep stage 1 (N1), iii. Non-REM sleep stage 2 (N2), and iv. Non-REM sleep stage 3 (N3). Any discrepancies in sleep staging (<5% of all epochs) were discussed in person with a board-certified sleep neurologist (M.M.L.) through adjudication sessions, and resolved upon consensus. PSG epochs were carefully examined for motion artifacts, which were used to eliminate potentially corrupt MRS data.

**Figure 2.**
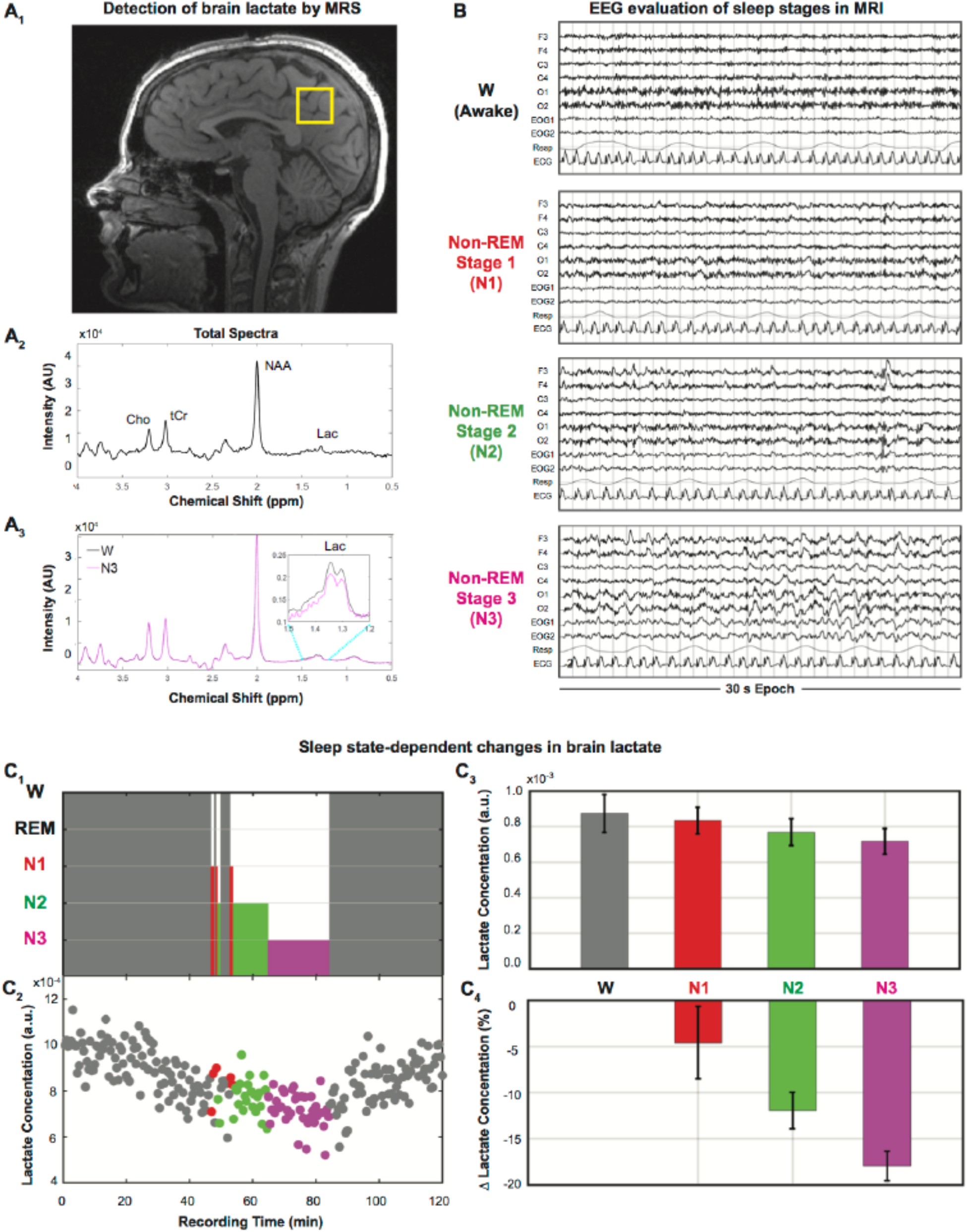
Lactate concentration in human brain is reduced during sleep relative to wake. **A_1_,** Measured single voxel (22 cm^3^, yellow square) on a sagittal T1-w MR image of a 27-year-old healthy young female subject. **A_2_**, A 7.5 sec 3T single voxel ^1^H-MR spectrum showing metabolites including lactate, **A_3_**, ^1^H MR spectra, averaged over wake and non-REM stage 3 sleep (N3) (intensity in normalized absolute units). Lactate concentrations are calculated as area of spectra within [1.23-1.48 ppm]. **B,** Representative sleep stages are visually scored for 30 s epochs. **C_1_**, Hypnogram and **C_2_,** corresponding lactate concentration for 30 s intervals. **C_3_,** Average lactate concentration in arbitrary units were: **0.87×10^-3^** during W (86 min), **0.83×10^-3^** during N1 (2.5 min),**0.76×10^-3^** during N2 (12 min), and **0.71×10^-3^** during N3 (19.5 min) present a decrease in sleep relative to awake with a decline of **C_4_,** 4.6 % during N1, 12.0 % during N2, and 18.0 % during N3 when compared to W.

#### MRI and ^1^H-MRS

Anatomical images were acquired using a T_2_-weighted fast spin echo (HASTE, repetition time (TR) 1500 ms, echo time (TE) 80 ms) and 3D T_1_-weighted gradient echo sequence (MPRAGE; TR 1900 ms; TE 2.52ms). For accurate non-invasive detection of brain lactate with high temporal resolution, we used a single voxel ^1^H-MRS technique that acquired signals from a fixed volume of interest (VOI) within a subject, with VOIs ranging from 12-24 cm^3^ across subjects with a focus on avoiding CSF spaces. See **Fig. 2A_1_** for a representative VOI position (Subject 1, 27 y female) located within the posterior cingulate cortex, a major node of the default network and area affected early in Alzheimer’s disease^25^. The ^1^H MRS protocol consisted of point-resolved spectroscopy sequence (PRESS) collected with TR 1875 ms, TE 270 ms and 4 signal averages with phase cycling, spectral width 1300 Hz, sampling points 1024, and total acquisition time ranged from 88 min to 180 min depending on the subject/experiment. Typical water line widths at full-width half maximum were between 12.2 - 14.3 Hz. MR spectra were collected every 7.5 s (**Fig. 2 A_2_**; Subject 1), which allowed the detection of several brain metabolites including choline (Cho), creatine (Cr), *N*-acetyl-aspartate (NAA), and lactate.

Three of the subjects (two females and one male; mean age 25.6 (± 2.8; SD) years; age range 24-29 years) returned for a follow-up visit to collect diffusion-sensitized PRESS MRS sequence for up to 180 min sleep/wake studies. For the diffusion sensitized MRS acquisition, diffusion weighting was achieved by simultaneously applying gradients along all three laboratory axes (δ=25 ms, Δ=36.5 ms, b=500 s/mm^2^) prior to and after the initial refocusing pulse of the PRESS sequence. The diffusion weighting was interleaved with no diffusion weighting at every TR value of 1875 ms, resulting in a complete pair (diffusion and nondiffusion MRS) being acquired every 3750 ms. For apparent diffusion coefficients (ADC) comparisons, MRS spectra from all sleep states (N1, N2, and N3) were averaged separately for diffusion weighted and non-diffusion weighted acquisitions.

Non-parametric analyses were performed to better appreciate how rapidly the lactate signal changed across sleep stages. This was accomplished using jMRUI ^26^ and MATLAB software packages [*R2017b; Mathworks, Natick, Mass*]. The MRS pre-prepossessing steps in jMRUI included post-acquisition water suppression, line broadening (Lorentzian, 5 Hz), automatic frequency shift and phase correction. Spectra were then imported to MATLAB to calculate the alterations in mean metabolite concentrations over the entire sleep-wake cycles. In order to determine and compare mean lactate signals for each sleep state across the recording duration, we averaged ^1^H MRS signals over a period of 30 s that coincided with every epoch of the PSG scoring. Lactate intensities were determined from non-parametric signal integrals over a defined spectral range [1.23 - 1.48 ppm], normalized to the peak area of nonsuppressed water signals [calculated over 4.3 - 5.1 ppm]. Processing of the ^1^H-MRS data was done in a blinded fashion with respect to the PSG data and motion artifacts based on PSG recordings were used to eliminate potentially corrupt MRS data for each subject. Mean lactate concentration in each stage (W, N1, N2, N3), as well as relative percent change in lactate concentration (%) from Wake to each sleep stage (N1, N2, N3) were calculated.

In order to examine group average arousal state dependent brain lactate concentration, we quantified mean lactate levels for each sleep stage across the nine subjects using LCModel.^27^ Each spectrum was zero-filled to 3072 points, line broadened (Lorentzian, 2.0 Hz) and frequency shifted to NAA peak and phase corrected. After digital water suppression by using HLSVD Propack filter (jMRUI-5.1 package),^28^ parametric fittings from LCModel were used to extract normalized signal intensities for lactate (Lac), total creatine (tCr), choline (Cho), and *N*-acetyl-aspartate (NAA). These concentration values were corrected for T_1_ and T_2_ relaxation effects.^29-31^ For NAA, tCr, Cho and Lac, T_1_ values of 1860, 1740, 1320 and 1340 ms; and T_2_ values of 262, 151, 199 and 239 ms were used respectively. A reference tCr metabolite concentration of 11.85 mM was assumed and signal ratios relative to tCr values obtained from LCModel fitting were corrected by dividing by transverse relaxation (exp(-TE/T_2_) and longitudinal relaxation (1-exp(-TR/T_1_)) factors. For diffusion-weighted PRESS data, spectra corresponding to wake and sleep stages (N1, N2, and N3) were binned and averaged separately into diffusion weighted and non-diffusion weighted scans after the described pre-processing steps. Water suppressed spectra were fit using LCModel software. The ADC (apparent diffusion coefficient, mm^2^/s) values for all metabolites (m) in different sleep stages were calculated using the fitted signal values from non-diffusion weighted MRS acquisitions (S0_m_) and diffusion weighted acquisitions (S1_m_) by using the following relation:

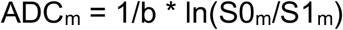

where b corresponds to a value of 500 s/mm^2^.

### Statistical Analysis

Statistical analysis of MRS data used linear mixed-effects models to assess the association between sleep stages and change in species concentration within subjects. The fixed-effects component used sleep staging as a four-level factor with the wake (W) state as the baseline reference while random-effects blocked on individual subjects with random intercepts to account for repeated measures and serial correlation. Prior to analysis, metabolite concentration outcomes were first normalized to the water signal at the individual subject level. Both absolute concentration and relative (fractional) differences were assessed, with models of fractional change including mean concentration at baseline as an additional covariate. No difference was observed between models using either outcome. Normality assumptions and model diagnostics evaluating influence and leverage used a combination of inspection of studentized residuals and formal fit criteria including Cook’s distance and the standardized difference of the betas. No outliers or excessive leverage points were identified for either specific mean concentration observations nor for specific subjects. Paired t-tests were used to compare metabolite ADC values between sleep and wake. All statistical analyses were conducted using R 3.5 ^32^ with additional utility using the lme4 package for linear mixed effects modeling.^33^

## Results

A representative hypnogram, depicting Subject 1’s sleep-wake cycles across the 120 min experiment time, is plotted with corresponding normalized lactate levels for each 30 s interval (**Fig. 2C_1-2_**) which demonstrates a decrease in brain lactate from baseline wakefulness (W: 50 min) through different sleep stages. To determine and compare mean lactate levels for each state across the duration of the recording, we averaged lactate levels for each arousal state (**Fig. 2C_3_**). Note that all wake stages were averaged regardless of pre-sleep or wake after sleep onset. For this subject, when compared to state W (86 min), normalized lactate levels were 4.6 % lower in N1 (2.5 min), 12.0 % lower in N2 (12 min), and 18.0 % lower in N3 (19.5 min) (**Fig. 2C_4_**). Averaged spectra from W and N3 sleep stages are plotted for a comparison with an inset showing the lactate doublet peak region (**Fig. 2A_3_**). Despite the brevity of N1 (brevity consistent with that expected in healthy subjects), lactate levels were still determined to be lower during N1 relative to W.

To examine group averaged, arousal state-dependent lactate concentration, we quantified mean lactate signal for each arousal state across the subjects using LCModel;^27^ a parametric spectral modeling software program. All nine subjects cycled through W (mean duration 48.8 ± 34.7 min) and N1 (mean duration: 9.1 ± 5.2 min); eight subjects cycled into N2 (mean duration: 36.5. ± 23.4 min, N=8), and six subjects cycled into N3 (mean duration: 19.3±13.2 min, N=6). Because only three of the subjects achieved REM sleep, data from their REM epochs were not included in the analyses. Cerebral metabolite concentration across each arousal state is summarized in **Table 1A**. The brain MRS lactate signal originates from all tissue spaces (intracellular, extracellular, blood) and across all subjects averaged 0.65 (0.14) mM in the wake (W) state, a value consistent with literature reports ^34^. When compared to W, average lactate concentrations within each sleep stage showed a reduction of [4.9 ± 4.9] % in N1, [10.4 ± 5.2] % in N2, and [24.0 ± 5.8] % in N3 (**Fig. 3, Table 1B**). These results are consistent with the previous rodent studies ^12-17^ also reporting reduced brain lactate levels during sleep when compared to wakefulness ranging from 12-35 %. Note that we did not observe significant changes in the ^1^H_2_O, Cho, Cr, or NAA signals across the time series or arousal levels (see **Table 2**).

**Figure 3.**
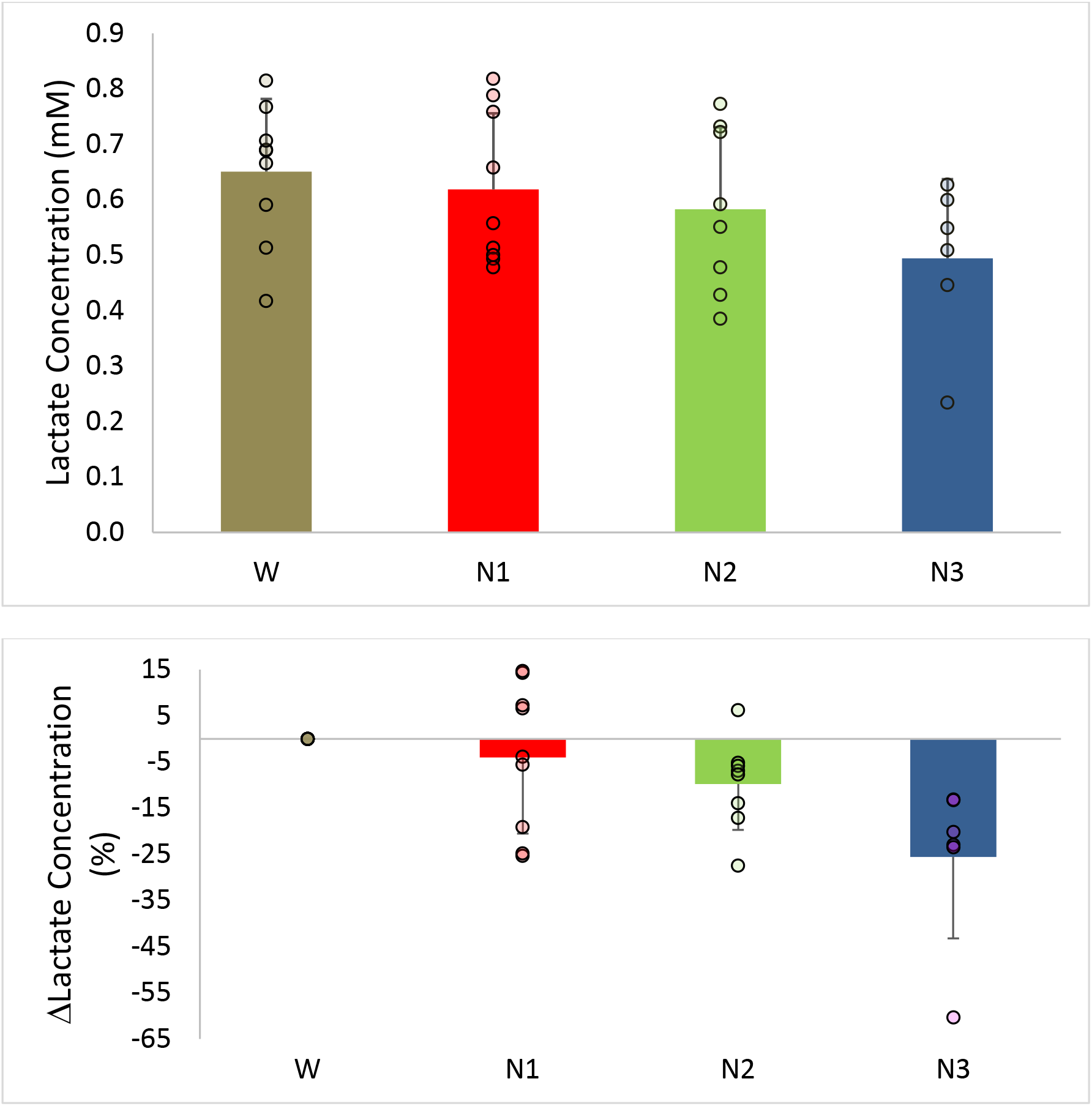
Group average brain lactate concentrations (N=9). **A.** Total lactate concentrations: **0.65 (0.05) mM,** mean (SD), during W (mean duration 48.8 ± 34.7 min, nine subjects), **0.62 (0.05) mM** during N1 (mean duration: 9.1 ± 5.2 min, nine subjects), **0.58 (0.05) mM** during N2 (mean duration: 36.5. ± 23.4 min, eight subjects), and **0.49 (0.06) mM** during N3 (mean duration: 19.3±13.2 min, six subjects). **B.** The change in lactate concentration between sleep states and awake; N1 not significantly different than W; N2 decreased 11% (P<0.05) compared to W; N3 decreased 24% compared to W (P<0.001). Individual measurements are indicated by circles.

**Table 1A.** Group Average Metabolite Concentration in Each Arousal State

| MRS Signal | Arousal State |  |  |  |
| --- | --- | --- | --- | --- |
|  | W | N1 | N2 | N3 |
| Lac | 0.65 (0.13) | 0.62 (0.14) | 0.58 (0.15)* | 0.49 (0.14)** |
| NAA | 16.26 (4.10) | 16.04 (4.19) | 15.65 (4.15) | 15.98 (3.63) |
| tCr | 11.85 (2.71) | 11.72 (2.81) | 11.41 (2.87) | 11.85 (2.42) |
| Cho | 1.95 (0.53) | 1.98 (0.56) | 1.90 (0.55) | 1.87 (0.31) |

Values are presented as mean (SD) with units of mM
\* P = 0.05; \*\* P < 0.001 for lactate decrease relative to W

**Table 1B.**
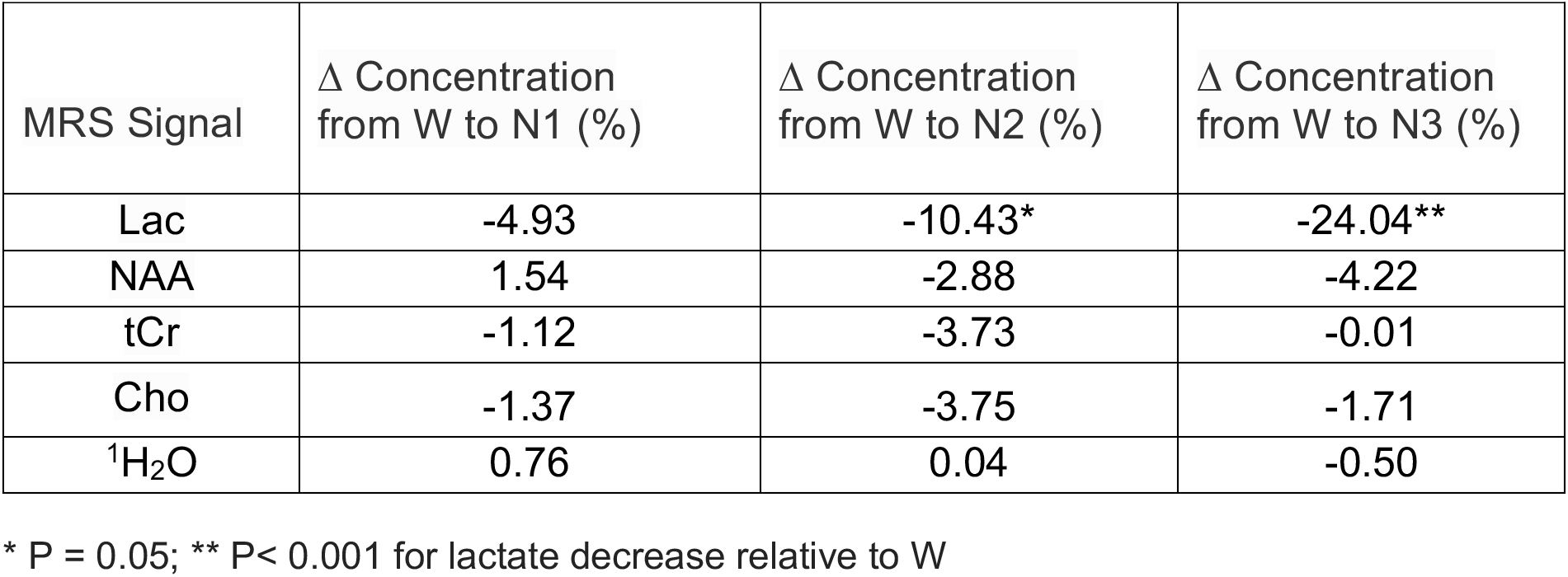
Group Average Changes in Metabolite Concentration

| MRS Signal | Δ Concentration from W to N1 (%) | Δ Concentration from W to N2 (%) | Δ Concentration from W to N3 (%) |
| --- | --- | --- | --- |
| Lac | -4.93 | -10.43* | -24.04** |
| NAA | 1.54 | -2.88 | -4.22 |
| tCr | -1.12 | -3.73 | -0.01 |
| Cho | -1.37 | -3.75 | -1.71 |
| <sup>1</sup>H<sub>2</sub>O | 0.76 | 0.04 | -0.50 |

\* P = 0.05; \*\* P < 0.001 for lactate decrease relative to W

**Table 2.** Group MRS ADC Values in Wake and Sleep

|  | Lactate | NAA | tCreatine | Choline | H<sub>2</sub>O |
| --- | --- | --- | --- | --- | --- |
| **Awake** | 1.32 (0.31) | 0.29 (0.08) | 0.24 (0.13) | 0.17 (0.11) | 1.80 (0.10) |
| **Sleep** | 0.50 (0.32)* | 0.27 (0.10) | 0.26 (0.03) | 0.21 (0.01) | 1.79 (0.12) |

Values are presented as mean (SD) with units of 10<sup>-3</sup> mm<sup>2</sup>/s
P < 0.002 for lactate ADC decrease relative to W

The ADC of lactate differs between intracellular and extracellular (interstitial) compartments, with lactate in the extracellular space reporting a substantially larger ADC relative to intracellular space.^35^ If the lactate signal decrease during sleep is associated with the selective loss of extracellular lactate, lactate diffusivity would be expected to differ markedly between wake and sleep. To investigate this, we quantified the ADC of lactate by averaging across wake (W) and combined sleep (N1, N2, N3) epochs (**Fig. 4**). The average time for wake state was 40 ± 19 (mean ±SD) min, and sleep was 96 ± 39 min. We observed a significant decrease in lactate ADC accompanied by a decreased brain lactate concentration in sleep compared to wake (P<0.002). There were no differences in ADC values between wake and sleep for H_2_O, NAA, tCr, or Cho.

**Figure 4.**
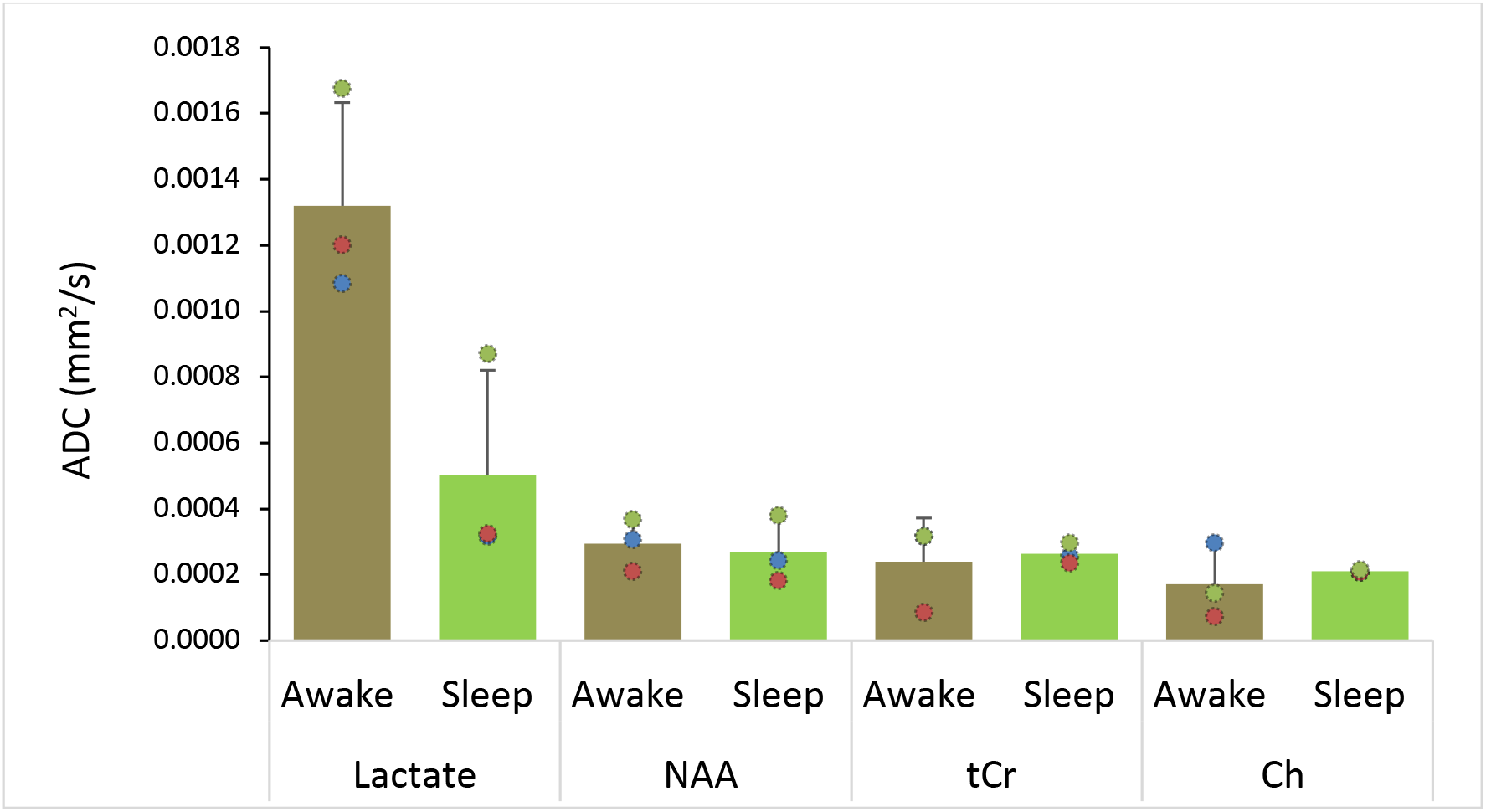
Metabolite Apparent Diffusion Coefficient (ADC) between sleep and wake states. Measurements from three subjects averaged for wake and sleep (N1, N2, and N3) states show marked reduction in lactate ADC in sleep compared to wake (P<0.002). ADC values in Wake were significantly greater for lactate than NAA, tCR, or Ch (P < 0.006). ADC values in Sleep were not different between lactate and NAA, tCr, or Ch (P > 0.18). No significant changes in ADC between wake and sleep states were observed for NAA, tCr, of Ch signals (P > 0.35). Mean values are plotted and error bars represent one standard deviation. Individual measurements are indicated by circles.

## Discussion

The combination of ^1^H MRS and PSG provided high quality quantifiable signals across several hours for each, and was safe and well-tolerated by all subjects. Substantial sleep durations in the scanner were realized by all of our study subjects with the majority of subjects experiencing prolonged N3 periods. The principal finding is a marked reduction in brain lactate concentration during sleep compared to wakefulness, with no change in NAA, tCr, or Cho signal levels. This observation is consistent with, and extends results from invasive small animal brain studies first reported more than 70 years ago.^12^ Additionally, we find a decrease in lactate ADC value that accompanied decreased brain lactate concentration in sleep compared to wake. Taken together, these findings suggest a sleep-dependent loss in brain lactate, predominantly from the extracellular pool, in agreement with prior small animal studies.^13,15^

There are three mechanisms that could explain decreased brain lactate levels during sleep relative to wakefulness: i) lactate production relative to consumption is reduced during sleep, ii) lactate consumption relative to production is increased during sleep, or iii) lactate clearance from the brain is increased during sleep. The first two mechanisms are metabolic in nature, and the third is physiologic and consistent with the studies of glymphatic function reporting increased CSF-ISF interchange during deep sleep. ^20,22^

First, we acknowledge the challenge and the opportunity of using MRS to quantify brain lactate. MRS provides a high-fidelity approach to quantify small molecules within tissue, and while present at high concentration within brain from a biochemical perspective, lactate concentration under normal conditions is on the lower end of MRS detectability. Our measurements had the advantages of MRS signal collection over extended times (2-3 hours total for 4 arousal states) and from a large volume of brain tissue with excellent magnetic field homogeneity. We find an average lactate concentration of 0.65 mM in the awake state, a value in good agreement with values reported from cerebral ^1^H MRS studies spanning three decades.^34,36^ Brain extracellular volume fractions are approximately 20% in the awake state ^37^ and average lactate concentration in the extracellular space is higher than the overall average brain lactate concentrations.^34,38,39^ During normal physiology in a wakeful state, the extracellular lactate pool may account for 50% of the MRS brain lactate signal, and offers the potential for detecting physiological changes associated with this space.^29,40^

Small animal studies report a marked loss of extracellular lactate during sleep.^13,15^ Our observation of decreased lactate ADC during sleep (**Fig. 3**), with values approaching that of other intracellular metabolites supports the interpretation of a selective loss of the more diffusible extracellular lactate, which would result in a decreased ADC due to a proportionately larger contribution of lactate signal from less diffusible intracellular lactate pools during sleep. No difference is ADC values were observed between wake and sleep for NAA, tCr, or Cho, which all are considered intracellular metabolites. No difference in ^1^H_2_O ADC was observed between wake and sleep; an observation supporting the notion that convective flow during sleep if increased, is increased to a small extent. ^41^ Brain lactate concentration change across sleep-wake cycles is not instantaneous, and in small animal studies a time constant of approximately 15 min is found.^15^ We observed similar kinetics for lactate concentration change across sleep-wake cycles in human brain. Since many of the subjects displayed some degree of sleep fragmentation while in the MR scanner, it is unlikely that our group average lactate concentrations represented true steady-state levels for any sleep state, which would have required prolonged (~45 min) continuity in each sleep state.

Glucose is the brain’s preferred energy source and is directly utilized by neurons and glia. Quantitative ^18^F-flourodeoxyglucose PET studies in young adults report large decreases in cerebral glucose utilization in deep sleep compared to wakeful states.^42^ Astrocytes support glucose uptake from plasma, convert some of this via aerobic glycolysis to lactate, which can be released into the interstitial space,^43,44^ or stored in a glycogen pool.^45,46^ The interstitial lactate may be taken up by neurons for aerobic energy production.^47^ Lactate may be oxidized in preference to glucose in neurons, particularly during times of intense neuronal activation, even when glucose is readily available.^48^ Extracellular brain lactate level decreased during SWS compared to wakefulness in freely behaving rats.^14^ Similar to natural sleep, anesthesia reduces astrocytic glycolysis and brain lactate concentration. ^49^ Interestingly, lactate concentration measured from cortical brain slices harvested from young adult rats in wake (dark) and sleep (light) periods, was reduced 25% in sleep compared to wake states^49^, and the administration of anesthetic agents dynamically reduced extracellular lactate despite a preparation that must disrupt key components important in glymphatic physiology. ^13,49^

Unlike muscle or liver, the brain does not possess large stores of glycogen to provide a reserve carbon source and extracellular lactate might partially fulfill this need. Glycogen stores within the brain (~7.8 umol/g brain about 1/4 that found for muscle ^50^) are localized primarily within astrocytes and these stores can serve as a reservoir for lactate.^46,51,52^

Astrocytic glycogen is depleted throughout the day and replenished during sleep,^45^ and it has been argued that glycogen restoration is a primary sleep function.^45^ During sleep or anesthesia, interstitial lactate may be taken up by astrocytes and converted back to glycogen; a metabolic pathway essentially running in reverse in sleep compared to wakeful states. Reduced cerebral blood flow^18,53^ and glucose utilization^42^ observed in sleep and during SWS in particular supports the notion that sleep associated lactate decline is at least partially due to decreased metabolic demand.

Brain lactate decline during sleep may also result from increased glymphatic activity.^20-22^ Lundgaard and colleagues^13^ used extracellular lactate as a surrogate marker for interstitial fluid clearance in brain, and argued that increased clearance plays a critical role in sleepwake state-dependent alterations brain lactate levels in mice. During periods of increased interstitial clearance associated with sleep or anesthesia, lactate levels in the cortex declined while lactate levels in deep cervical lymph nodes increased, suggesting that interstitial lactate is transported out of the brain, at least partially, via glymphatic-lymphatic coupling.^13^ While our finding that brain H_2_O ADC did not differ between sleep and wake argues against more rapid interstitial convective flow and glymphatic function during sleep, changes in other elements of water diffusion (e.g. metabolically driven transcytolemmal water flux ^54^) could potentially obscure the sleep-wake effects on slow interstitial convection. Current estimates of interstitial convective flow based on measurements made on the microscopic scale are low, and it is predicted that for small molecules such as lactate (molecular weight 90 g/mol) local thermal diffusion dominates the influence of convection.^37,41,55^ A recent extended time-series (24-48 h) analysis of contrast agent distribution across the human brain cortex, following intrathecal injection, estimated a larger gadobutrol (558 g/mol) ADC than ^1^H_2_O ADC measured by conventional (instantaneous) diffusion tensor imaging.^56^ This indicates that paravascular and perhaps interstitial convective flows, particularly across the large anatomical structures of the human brain, remain incompletely understood and warrants further investigation.^57^

### Limitations

Despite the novelty of the measurements and results presented above, there were limitations associated with this study. Only a single anatomical location was investigated and it is unclear how generalizable our findings are to the whole brain. The measured metabolite signal levels represent volume weighted averages from intracellular, extracellular, and even blood spaces (albeit quite small weighting), so a clear picture of changes within specific spaces is not directly available. ADC values reported here were determined from two b-values at long echo-times, and while trends are likely valid, ADC accuracy based on a two-point calculation is a concern, especially for low intensity signals such as lactate. Baseline brain lactate concentration during wakefulness could be affected by an overnight sleep deprivation, but this may be unlikely since others ^58^ have reported that 40-hours of sleep deprivation does not change baseline brain lactate concentration in either young (19-24 y) or older (60-68 y) healthy subjects while awake. Due to vendor compatibility with MRI, the PSG measurements did not include EMG, and instead EMG was substituted with a 3D accelerometer Finally, the number of subjects in the study was limited and a larger study needs to be performed to replicate the findings presented.

In conclusion, we have developed a novel non-invasive approach combining synchronized ^1^H MRS and PSG to investigate metabolic differences across sleep-wake cycles in human brain. Our observation of reduced lactate concentration during sleep, together with reduced lactate ADC, suggests a selective loss of the extracellular lactate pool. The merits of the metabolic and glymphatic mechanisms to explain decreased lactate concentration during sleep were considered, and it should be appreciated that these mechanisms are not mutually exclusive. Further studies are required to elucidate mechanistic details associated with changes in brain lactate concentration and dynamics during sleep. Nevertheless, the findings from the lactate measurements across sleep-wake cycles in human brain presented here provide a novel non-invasive approach to investigate fundamental aspects of sleep physiology.

## Author contributions

W.D.R., M.M.L., and J.J.I. obtained funding and conceptualized overall study design. W.D.R and M.K.S. developed the magnetic resonance spectroscopy (MRS) techniques. S.Y., M.M.L., and K.P. developed the polysomnography data collection, processing and analysis protocol. S.Y., K.P., M.K.S., and W.D.R carried out the experiments. S.Y. and M.K.S. processed the MRS data. S.Y. performed the non-parametric and M.K.S. preformed the parametric MRS data analysis. S.Y. and K.P. scored sleep stages and M.M.L. supervised. C.M. assisted in the statistical analysis. S.Y. and W.D.R wrote the manuscript, and all authors edited the manuscript.

## Acknowledgments

We thank study participants for their participation in the study, senior MR technologist William Woodward for conducting the MR scans, Brain Products GmBH and BrainVision for providing technical assistance, and numerous staff and collogues for their support of the study.

